# Super-resolution microscopy unveils FIP200-scaffolded, cup-shaped organization of mammalian autophagic initiation machinery

**DOI:** 10.1101/712828

**Authors:** Samuel J Kenny, Xuyan (Shirley) Chen, Liang Ge, Ke Xu

**Affiliations:** Department of Chemistry, University of California, Berkeley, CA 94720, USA; Chan Zuckerberg Biohub, San Francisco, CA 94158, USA; School of Life Sciences, Tsinghua University, Beijing 100084, China

## Abstract

Autophagy is an essential physiological process by which eukaryotic cells degrade and recycle cellular materials. Although the biochemical hierarchies of the mammalian autophagy pathway have been identified, questions remain regarding the sequence, subcellular location, and structural requirements of autophagosome formation. Here, we characterize the structural organization of key components of the mammalian autophagic initiation machinery at ∼20 nm spatial resolution via three-color, three-dimensional super-resolution fluorescence microscopy. We thus show that upon cell starvation, FIP200, a large structural protein of the ULK1 complex with no direct yeast homolog, scaffolds the formation of cup-like structures located at SEC12-enriched remodeled ER-exit sites prior to LC3 lipidation. This cup scaffold, then, provides a structural asymmetry to enforce the directional recruitment of downstream components, including the Atg12-Atg5-Atg16 complex, WIPI2, and LC3, to the cup inside. Moreover, we provide evidence that the early autophagic machinery is recruited in its entirety to these cup structures prior to LC3 lipidation, and gradually disperses and dissociates on the outer face of the phagophore membrane during elongation. We thus shed new light on the physical process of mammalian autophagic initiation and development at the nanometer-scale.

## Introduction

Macroautophagy, hereafter autophagy, is an essential catabolic process by which eukaryotic cells degrade and recycle organelles and proteins^1-8^. The general mechanism of autophagy begins with the activation of the ULK1 complex by mTORC1 in response to stresses. Subsequent recruitment and activation of the PI3K complex, PtdIns3*P* effectors (WIPI proteins), and the Atg12-Atg5-Atg16 complex enable the lipidation of LC3, which drives the formation of a cup-shaped membrane structure known as the isolation membrane, or phagophore. The phagophore engulfs cargo and closes before fusing with lysosome for cargo degradation. Together, the phagophore membrane with its associated protein machinery constitutes the autophagosome.

Although the biochemical hierarchy of autophagic machinery has been studied extensively^1,4^, open questions remain, especially regarding the spatiotemporal dynamics of autophagosome formation^6-8^. In particular, although the overall mechanism of autophagy is conserved across eukaryotes, autophagy initiation in yeast occurs at a single location (the pre-autophagosomal structure) whereas mammalian autophagosomes form throughout the cytosol, likely scaffolded by the ER or related tubulovesicular networks^4,9-11^. Meanwhile, many proteins essential for autophagy in mammals have no direct homologs in yeasts^1,3,7^. Understanding the functions of these proteins, particularly regarding their role in the structural organization of early autophagosomes, is of critical importance.

FIP200, also known as RB1CC1, is a component of the mammalian ULK1 complex, and may be involved in the spatial targeting and scaffolding of early autophagosomes^9,12^. Although FIP200 is a functional counterpart of yeast Atg17, the two share little sequence homology, but are instead related by a shared biochemical role as members of the ULK1 complex, and by their predicted flexible helical structure^9,10,13-15^, which may help scaffold the initial structure of autophagosomes^12,16^. As a SEC12 interactor, FIP200 further provides a possible link between the early autophagic machinery and ER-exit sites (ERES)^17^. Moreover, its proposed ability to recruit downstream components, including direct interaction with Atg16^18^ and tethering of Atg9 vesicles via Atg13^7,19^, provides mechanisms for the sequestration of a membrane source and subsequent LC3 lipidation. Whereas the biochemical interactions of FIP200 have been examined in depth, how exactly FIP200 acts as a structural scaffold during autophagosome formation remains speculative^12^. Elucidating this role with respect to the spatiotemporal dynamics of autophagosome development requires a detailed view of its organization into the nanometer size scale of nascent autophagosomes.

Recent advances in super-resolution fluorescence microscopy^20,21^, including three-dimensional (3D) stochastic optical reconstruction microscopy (STORM)^22,23^, provide new possibilities to achieve ∼20 nm spatial resolution and excellent target specificity with minimal invasion. We have recently used three-color 3D-STORM to identify a new mechanism for the remodeling of ERES as a prelude to autophagosome biogenesis^17^, and previously employed two-color 3D-STORM to visualize autophagy-mediated secretion^24^. Meanwhile, Karanasios *et al*. have used two-color two-dimensional (2D) STORM to examine the ULK1 complex in autophagy initiation^11^. Structured illumination microscopy has also been used to achieve ∼120 nm resolution in autophagy studies^11,25-27^.

Here, we use three-color 3D-STORM to reveal the ultrastructural organization of multiple protein targets during autophagosome development, hence connecting key biochemical complexes to spatial organizations at the nanometer-scale.

## Results

We started by developing a protocol for the preservation and STORM of LC3-positive phagophore membranes in mammalian cells. Induction of autophagy by starvation of U2OS cells in an amino-acid-depleted buffer, followed by a fixation and immunofluorescence protocol optimized for the preservation of fragile membrane structures (Methods), resulted in the observation of numerous diffraction-limited LC3 puncta under conventional epifluorescence microscopy (Fig. 1A inset).

**Figure 1.**
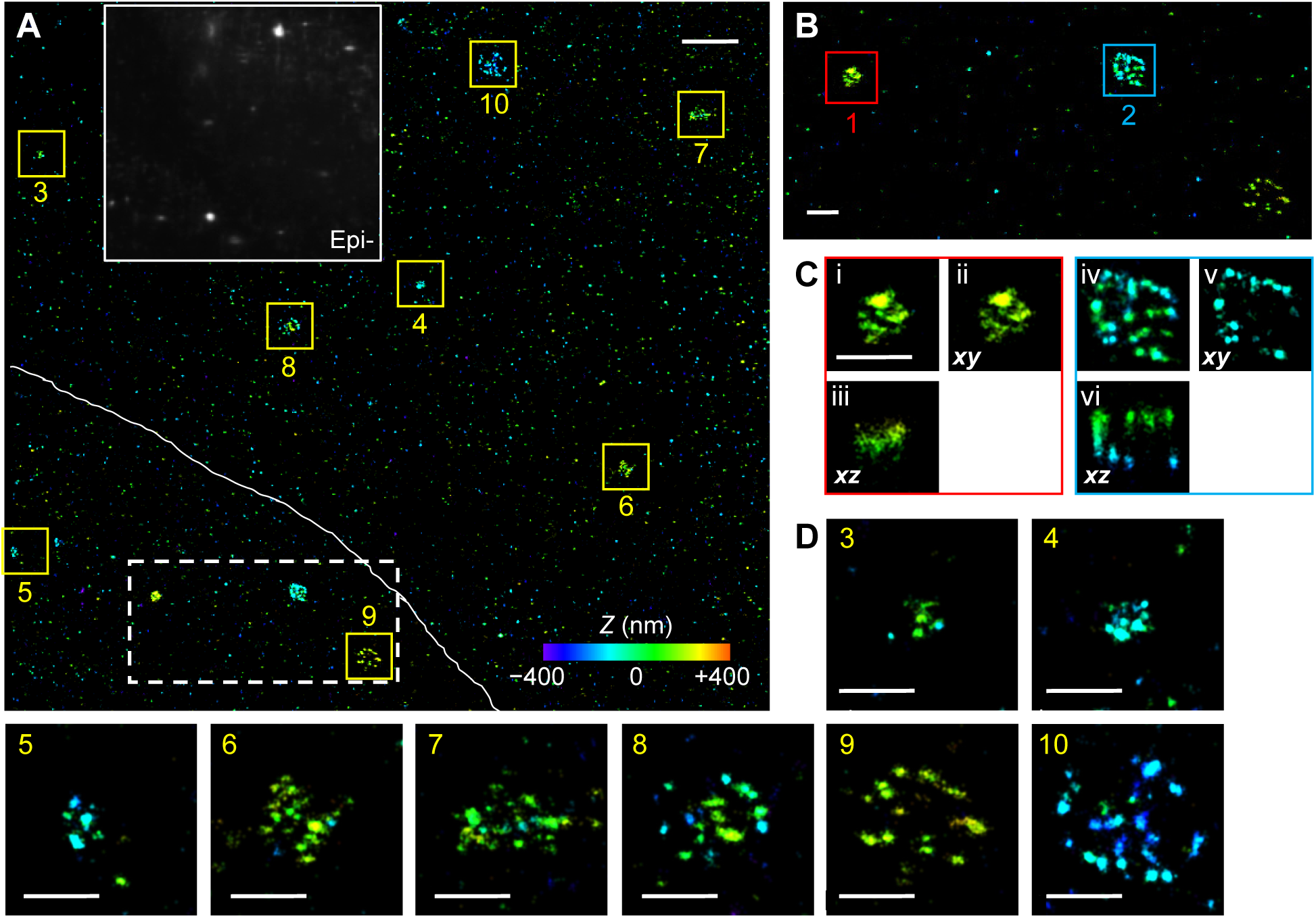
3D-STORM unveils diverse LC3 structures at the nanometer-scale. (A) 3D-STORM image of immunolabeled LC3 in starved U2OS cells. Depth (*Z*) position is color-coded (color scale bar). White line: boundary between two cells. Inset: diffraction-limited epifluorescence image of the same view. (B) Zoom-in of the area enclosed by the white box, containing three representative structures. (C) Further zoom-in of two structures 1 and 2 in (B), corresponding to early (red box) and intermediate (blue box) autophagosomes, respectively. (i,iv) Full *XY*-projections; (ii,v) Virtual sections in the *X*-*Y* plane for the central 150 nm depth in *Z*; (iii,vi) Virtual cross-sections in the *X*-*Z* plane for central slices 150-nm thick in Y. (D) Zoom-in for the structures 3-10 marked by the yellow boxes in (A). Scale bars: 2 μm (A), 500 nm (B-D).

3D-STORM super-resolution microscopy revealed that, at the nanometer-scale, the diffraction-limited LC3 puncta exhibited a variety of structural forms (Fig. 1A and Fig. S1). Notably, we found the smallest (<∼250 nm) LC3 structures, presumably in the early stage of autophagy, to be small patches (Fig. 1BC), whereas the medium (250-500 nm) structures were hemispherical cups (Fig. 1BC), and the largest (500-2000 nm), mature structures were enclosed spherical vesicles (Fig. 1D). The outstanding 3D capability of 3D-STORM was instrumental in identifying the correct geometry. For example, whereas the in-plane 2D projection in Fig. 1C (iv) suggests a spherical vesicle, virtual sections of the 3D data in the X-Y plane for the central depth Z (v) and in the X-Z plane for the central slice of Y (vi) showed that the structure was actually a hemispherical cup.

We next performed two-color 3D-STORM for FIP200 and LC3. This unveiled that FIP200 formed asymmetrically positioned, densely-packed cups around early LC3-positive phagophores (Fig. 2A, red arrow, and Fig. S2), but was largely absent from the large, spherical LC3 structures (Fig. 2A, blue arrow).

**Figure 2.**
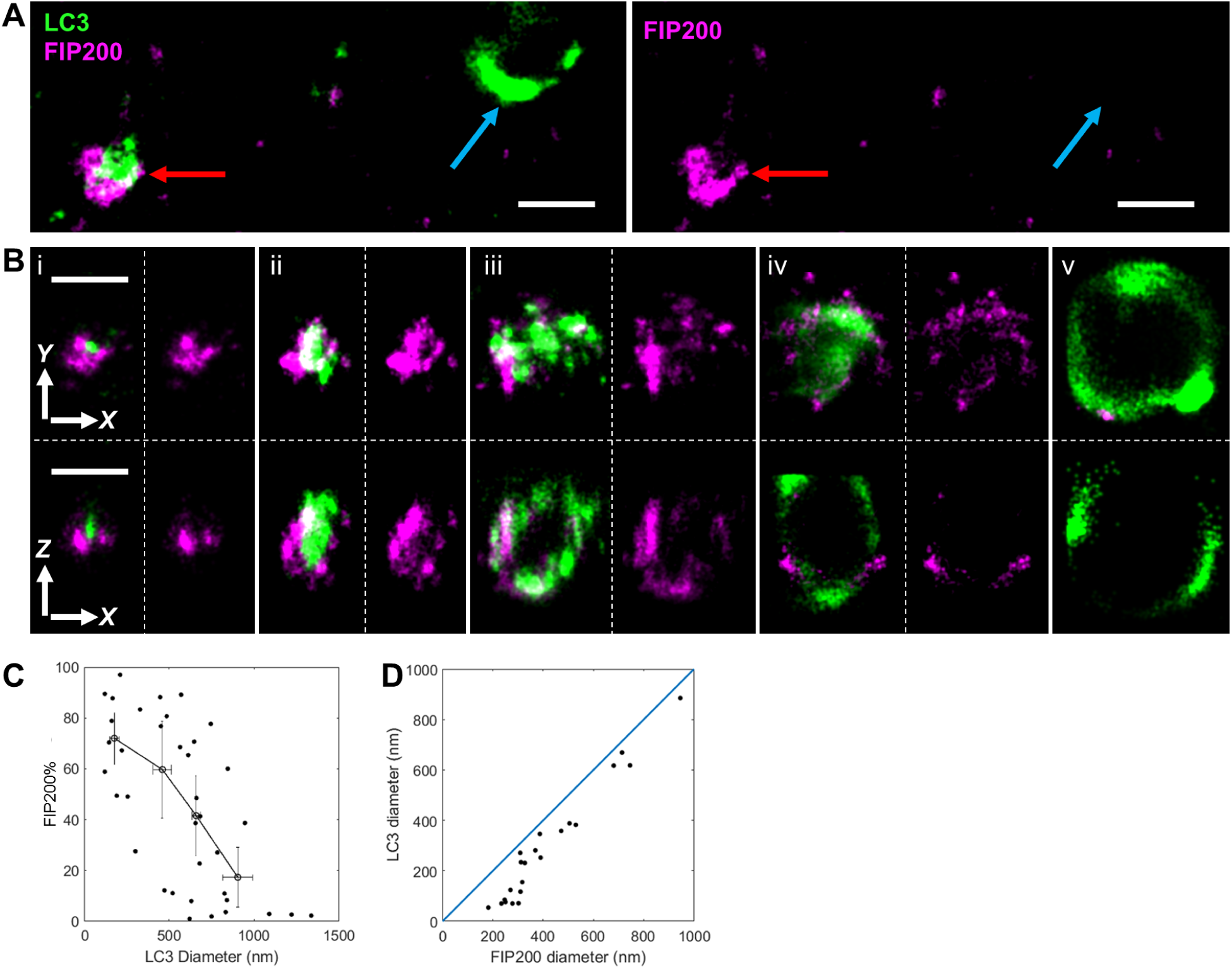
Two-color 3D-STORM of immunolabeled FIP200 (magenta) and LC3 (green) in starved U2OS cells. (A) Two structures in the same view, showing FIP200 as a cup-structure around a small LC3-positive autophagosome (red arrow), but absent from a large LC3 structure (blue arrow). Left: merged two-color image; Right: the FIP200 channel alone. (B) Representative images of FIP200 and LC3 in order of increasing LC3 diameter, hence a putative time series of autophagosome development. For each structure, top and bottom panels show full projections in the X-Y plane and virtual cross-sections in the X-Z plane for central slices 150-nm thick in Y, respectively. For (i-iv), left and right panels show merged two-color images and the FIP200 channel alone, respectively. (v) shows the two-color image only. (C) Quantification of FIP200 as a percentage of the total 3D-STORM signal in two-color images with LC3 for autophagosomes of different sizes (*n* = 37). Data are binned into quartiles according to the LC3 diameter, with standard deviations in each quartile denoted by crosses. (D) The FIP200 and LC3 sizes determined from two-color 3D-STORM of 22 colocalized structures. Blue line marks a slope of 1 (equal size). Scale bars: 500 nm (A-B).

To elucidate the progression of FIP200’s role in autophagosome development, we imaged structures of varying sizes and used the LC3 diameter as an indicator of developmental progression. The densely-packed cup-shaped motif of FIP200 was consistently observed around small (<∼250 nm) LC3 structures (Fig. 2B, i-ii; Fig. S2). The size of the smallest FIP200 cup structures that localized around single LC3 puncta (∼50 nm in size) was 260 ± 44 nm (*n* = 9; Fig. 2B, i and Fig. S2). The initially compact FIP200 labeling spread during phagophore elongation (Fig. 2B, ii and iii), and became sparsely distributed puncta decorating LC3-positive structures 250-500 nm in size (Fig. 2B, iii and iv). Minimal FIP200 signal was detected for large (>∼600 nm) LC3 structures (Fig. 2B, iv and Fig. 2A, blue arrow).

Quantification of the FIP200 signal as a percentage of the total 2-color STORM signal showed a monotonically decrease with increasing phagophore size (Fig. 2C). Comparison of the measured diameters of LC3 and FIP200 in colocalized structures (Fig. 2D) further showed FIP200 to be consistently larger. Together, our results indicate that FIP200 forms asymmetrically positioned cups around early autophagosomes, and remains at the outer face of the phagophore membrane while progressively dispersing and dissociating as the autophagosome develops.

The FIP200 cups we found above for early-stage phagophores are reminiscent of the cup-shaped SEC12 structures we recently reported in the FIP200-facilitated, starvation-induced remodeling of ERES^17^. Notably, three-color 3D-STORM showed that SEC12 colocalized well with FIP200 cups ∼200 nm in size in the absence of LC3 (Fig. 3A, i). Subsequent LC3 recruitment and phagophore elongation resulted in a quick drop in SEC12 signal (Fig. 3A, ii-iv), with minimal SEC12 observed for structures >∼400 nm in size, at which stages FIP200 was still present at high levels (Fig. 3A, iii-iv). These results indicate that the recruitment of FIP200 to the SEC12-enriched remodeled ERES and the formation of the observed cup-like motif occur prior to LC3 lipidation.

**Figure 3.**
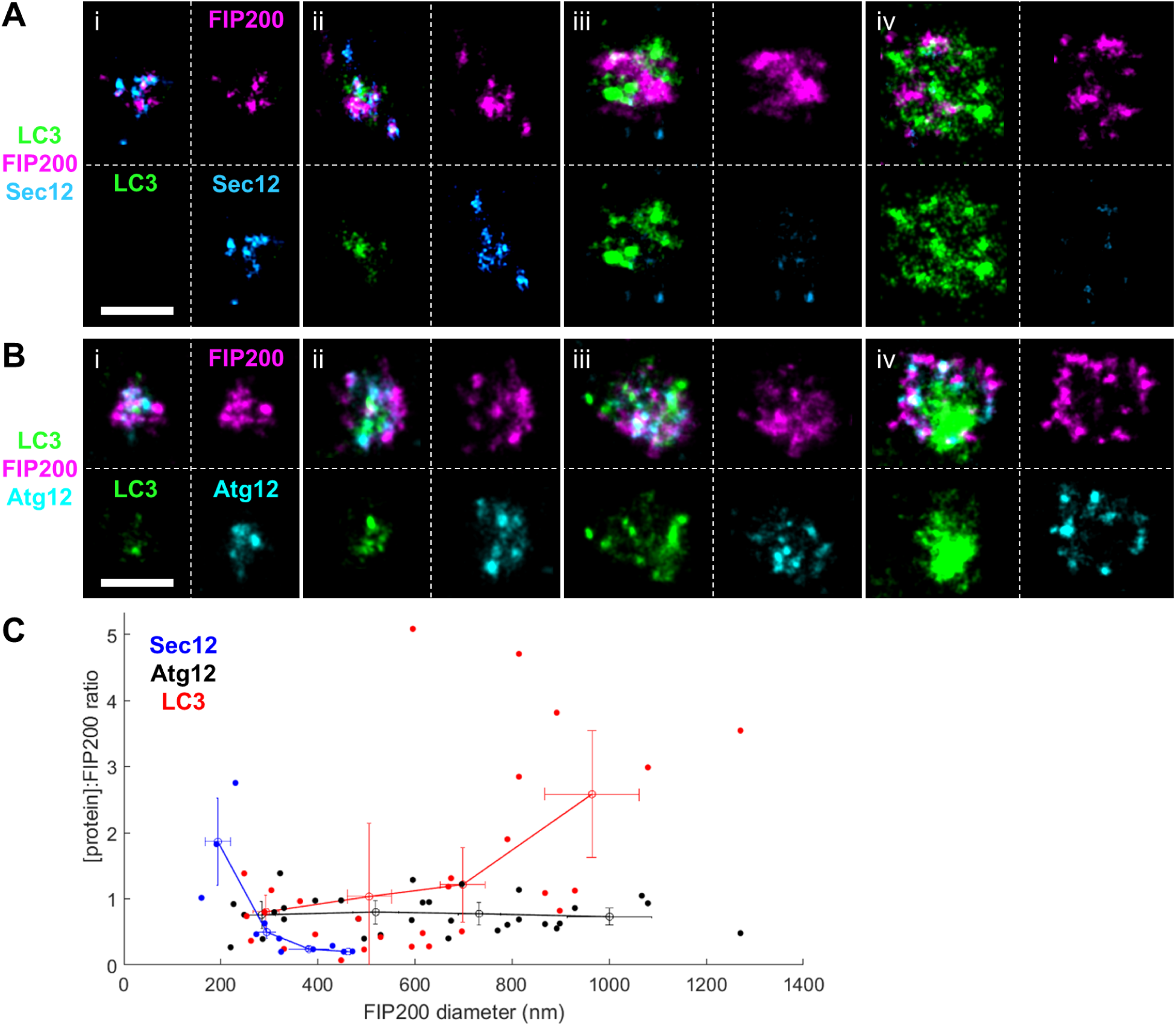
Three-color 3D-STORM indicates that FIP200 cups form at SEC12-enriched remodeled ERES, and define the structural organization of downstream components. (A) Representative three-color 3D-STORM images (merged 3-color data and separated color channels) of FIP200 (magenta), SEC12 (light blue), and LC3 (green) in early- and intermediate-stage autophagosomes. (i) A colocalized FIP200/SEC12 cup structure prior to LC3 lipidation. (ii-iv) SEC12 begins to dissociate upon LC3 lipidation (ii), and is absent from larger autophagosomes (iii,iv). (B) Representative three-color 3D-STORM images (merged 3-color data and separated color channels) of FIP200 (magenta), Atg12 (cyan), and LC3 (green) at similar stages. (i) Atg12 recruitment to the FIP200 cup at or before LC3 lipidation. (ii-iv) Atg12 matches the localization pattern of FIP200 during phagophore elongation. (C) Calculated protein count ratio between FIP200 and SEC12 (*n* = 11; blue), Atg12 (*n* = 31; black), or LC3 (*n* = 27; red) for colocalized structures. Lines represent data binned into quartiles according to the FIP200 diameter, with standard error for each quartile denoted by crosses. Scale bars: 500 nm (A,B).

We next examined how the unique FIP200 cup structure recruits downstream components. The Atg12-Atg5-Atg16 complex is the last major protein complex to be recruited prior to LC3 lipidation and phagophore elongation^6^. As FIP200 directly interacts with the middle region of Atg16^18^, it provides a link between the Atg12-Atg5-Atg16 and ULK1 complexes. Remarkably, three-color 3D-STORM showed that Atg12 colocalized closely with FIP200, and this agreement was observed across all stages of autophagosome development (Fig. 3B). Comparison of the individual color channels showed that within a single phagophore, many nanoscale FIP200 puncta have a corresponding Atg12 punctum (Fig. 3B). In comparison, the LC3 signal exhibited a range of total intensity with little direct correspondence to either the FIP200 or Atg12 signals (Fig. 2AB and Fig. 3B).

With the above data, we plotted the STORM signals of the SEC12, Atg12, and LC3 labeling at FIP200 cup structures as a function of structure size (Fig. 3C). This showed that SEC12 primarily localized to structures <250 nm in size, with the SEC12:FIP200 ratio rapidly falling off at the earliest stages of development. Meanwhile, Atg12 localized to FIP200 structures in a near 1:1 ratio independent of the structure size, whereas the LC3:FIP200 ratio increased quickly during phagophore elongation (300-800 nm).

Zoom-in of the three-color STORM images further revealed the relative spatial organization of different autophagic components. We thus found that FIP200 defines the outermost edge of autophagosomes, Atg12 localized to the inner edge of the FIP200 shell, whereas LC3 constituted the innermost part of the structures (Fig. 4A). We also examined WIPI2, a PtdIns3*P* effector that controls LC3 lipidation by interacting with Atg16L^28^: we found it localized inside the FIP200 cups (Fig. S3), and was at the innermost part relative to FIP200 and Atg12 in hemisphere structures (Fig. 4B), similar to what we observed for LC3.

**Figure 4.**
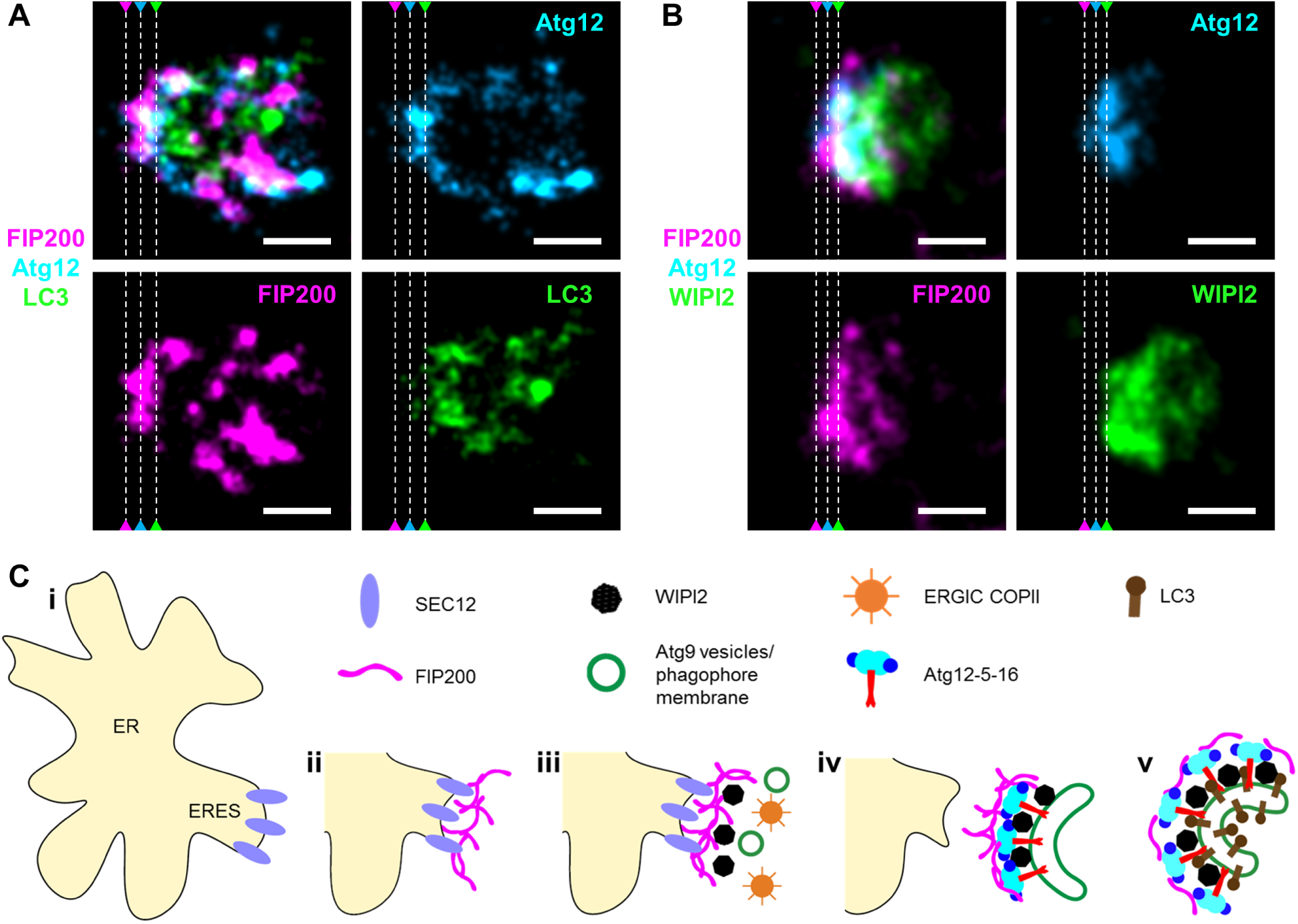
FIP200 provides a structural basis for the directional recruitment of downstream autophagic machinery. (A) Zoom-in of three-color STORM image of FIP200 (magenta), Atg12 (cyan) and LC3 (green) in a hemispherical cup structure, shown as merged and separated color channels. The three dotted lines mark the apparent edges of the three channels. (B) Three-color STORM image of FIP200 (magenta), Atg12 (cyan), and WIPI2 (green). The three dotted lines mark the apparent edges of the three channels. (C) Proposed model of phagophore scaffolding and elongation. (i) SEC12 clusters to ERES upon starvation. (ii) FIP200 recruitment to SEC12 forms dense cup-shaped scaffolds. (iii) Together with other members of the ULK1 complex, this scaffold recruits Atg9-tethered vesicles, COPII vesicles, and WIPI2. (iv) Atg12-5-16 is recruited to the inside of the FIP200 cup by interacting with both FIP200 and WIPI2 to facilitate vesicular fusion into a cup-shaped phagophore. (v) Lipidation of LC3 by Atg12-5-16 and other machineries drives phagophore elongation, during which process FIP200 and Atg12-5-16 remain at the outer face of the phagophore while dispersing and dissociating. Scale bars: 200 nm (A,B).

## Discussion

The signaling cascades and biochemical components required for autophagy initiation have been studied extensively^1,6,7^. While our understanding of mammalian autophagy has benefited greatly from comparison to yeast, significant differences exist between the two: in particular, mammalian autophagy occurs throughout the cytosol, rather than from a single initiation site. Mammalian proteins with no direct yeast homologs are possible candidates for these differences in spatial organization. FIP200 is one such protein: while functionally related to yeast Atg17^9,15^, it is much larger (200 kDa vs. 47 kDa), and as such, may play a unique role in scaffolding the mammalian autophagosome biogenesis. Additionally, its established role as a SEC12 interactor^17^, combined with the potential of the starvation-remodeled ERES as a site of autophagosome formation^11,17^, make it a compelling target of study.

Here, through three-color 3D-STORM, we found that in amino acid-starved cells, FIP200 was recruited to the SEC12-enriched remodeled ERES, where it formed cup-shaped autophagic precursors. The formation of such FIP200 cup structures may be a prerequisite for the subsequent recruitment of the autophagic machinery, and we found the smallest cup structures that colocalized with LC3 puncta to be ∼260 nm in size.

Since cups are inherently asymmetric, they provide a mechanism for the asymmetric elongation of the phagophore membrane. Three-color 3D-STORM directly visualized such asymmetry, with Atg12 primarily localized to the inside of FIP200, and WIPI2 and LC3 to the further inside of Atg12 (Fig. 4AB). The persistence of this directional recruitment implies an important role for the initial structural organization of the FIP200 scaffold in organizing the cup-shaped phagophore double membrane. The close colocalization and near 1:1 presence of Atg12 and FIP200 across different stages (Fig. 3B, C) further suggest that components upstream of LC3 are fully recruited to the cup structure prior to LC3 lipidation. The FIP200 cup-scaffolded mechanism also helps explain the general observation that Atg proteins tend to be recruited to the outer face of the phagophore membrane^29,30^, from where they then dissociate during autophagosome formation^25^.

As we observed dense FIP200 cups at early autophagosomes that colocalized with SEC12, it raises the question of whether SEC12 forms cup structures prior to FIP200 recruitment. Examination of our STORM data showed little evidence of SEC12 cups without FIP200 colocalization. Indeed, we have previously shown that the knock-down of FIP200 abolishes the enlargement of SEC12-ERES in starved cells^17^. In our recent work on SEC12 accumulation at ERES during the generation of COPII-coated vesicles in un-starved cells^31^, we find that SEC12 organizes into dense, flat sheets, as opposed to cups. These observations suggest that FIP200 provides a mechanism for the restructuring of the SEC12-enriched ERES membrane in starved cells towards cup formation. Clues as to FIP200’s ability to remodel membranes may be gleaned by recent simulations showing that the association of Atg17 dimers with tethered vesicles is sufficient to reshape these membranes into tube, disk, or cup shapes^16^.

In summary, we have shown that FIP200 defines a cup-shaped scaffold for the directional recruitment of subsequent autophagic machineries and hence the structure of nascent autophagosomes. Upon starvation, FIP200 is recruited to the SEC12-enriched, remodeled ERES, where it nucleates into a dense, cup-like scaffold (Fig. 4C, i-ii). This scaffold recruits other members of the ULK1 complex, which in turn recruits Atg9-tethered vesicles, as well as activates downstream PI3K complexes and PtdIns3*P* effectors. PI3K-dependent ERGIC-COPII vesicles^32^ and WIPI2 are hence recruited to this initiation site (Fig. 4C, iii). Subsequent recruitment of the Atg12-5-16 complex facilitates vesicular fusion into a cup-shaped phagophore double membrane (Fig. 4C, iv). Elongation of this membrane then proceeds with LC3 lipidation, during which process the *a priori* recruited ULK1 and Atg12-5-16 complexes gradually disperse and dissociate on the outer face of the phagophore membrane (Fig. 4C, v).

Our work also highlights how state-of-the-art multicolor super-resolution microscopy can help elucidate the intricate autophagy pathway. With ∼20 nm resolution, we both resolved well-defined structures for multiple diffraction-limited targets with their relative positions determined, and generated pseudo time sequences using the measured autophagosome size. Future live-cell super-resolution microscopy^20,21,33^ experiments, although technically highly demanding, may directly visualize these processes within individual structures, circumventing the need for examining many fixed snapshots.

## Acknowledgment

We thank Prof. James Hurley for discussion, and acknowledge support from the National Science Foundation under CHE-1554717 and the Pew Biomedical Scholars Award. K.X. is a Chan Zuckerberg Biohub investigator.

## Methods

### Cell culture and immunolabeling

U2OS cells (human osteosarcoma) were obtained from the UC-Berkeley cell culture facility. Cells were maintained in Dulbecco’s Modified Eagle Media plus 10% fetal bovine serum. Cells were cultured on #1.5 coverslips and starved by incubation in Earle’s Balanced Salt Solution for 1 h at 37 °C immediately before fixation. Cells were fixed in 4% paraformaldehyde in Dulbecco’s Phosphate-Buffered Saline (PBS) for 20 min at room temperature, washed three times with PBS, and then permeabilized with 40 μg/mL digitonin (Sigma D141) in PBS for 10 min. After blocking in a blocking buffer (BB; 3% w/v bovine serum albumin in PBS), samples were incubated with primary antibodies (below) in BB overnight at 4 °C and then washed three times with a washing buffer (0.1% w/v bovine serum albumin in PBS). Cells were incubated in dye-labeled secondary antibodies (below) in BB for 1 hour at room temperature, and then washed again with the washing buffer and PBS prior to imaging.

Rat anti-SEC12 antibodies (1:50) were purified from hybridoma clone 6B3 originally provided by the Kota Saito lab (University of Tokyo, Tokyo, Japan)^34^. The following commercial primary antibodies were used: mouse IgG1 anti-LC3 (MBL #M152-3; 1:100), rabbit anti-FIP200 (Proteintech #17250-1-AP; 1:100), mouse IgG2b anti-Atg12 (Genetex GTX629815; 1:100), and mouse IgG1 anti-WIPI2 (Sigma-Aldrich #MABC91; 1:100). Alexa Fluor 647-labeled secondary antibodies were from Invitrogen. CF680- and CF568-labeled secondary antibodies were prepared through reactions of the corresponding succinimidyl esters (Biotium) with secondary antibodies from Jackson ImmunoResearch.

### STORM imaging

3D-STORM imaging was performed on a homebuilt setup^35^ using a Nikon CFI Plan Apo λ 100x oil-immersion objective (NA 1.45), as described previously^17^. Briefly, the above dye-labeled cell samples were mounted with a Tris-HCl-based imaging buffer containing 5% (w/v) glucose, 100 mM cysteamine, 0.8 mg/mL glucose oxidase, and 40 µg/mL catalase^22,23^. The sample was illuminated by 647- or 560-nm lasers at ∼2 kW cm^-2^, which reached the sample at incident angles slightly smaller than the critical angle, thus illuminating a few micrometers into the sample. The resultant stochastic photoswitching of single-molecule fluorescence, which was further assisted by a weak (0-1 W cm^-2^) 405-nm laser, was recorded using an EM-CCD (Andor iXon Ultra 897) at 110 frames per second, for a total of ∼60,000 frames per image. A cylindrical lens was inserted into the imaging path to encode the depth (z) information into the single-molecule image shape^23^. The raw STORM data were analyzed according to previously described methods^22,23^.

Two-color 3D-STORM imaging was performed for targets labeled by Alexa Fluor 647 and CF680. With 647-nm excitation, a ratiometric detection scheme^36,37^ was employed to concurrently collect the emission of Alexa Fluor 647 and CF680 single molecules. Emission of these dyes was split into two light paths using a long pass dichroic mirror (T685lpxr; Chroma), each of which was projected onto one half of the EM-CCD camera. Dye assignment was performed by comparing the intensity of each single molecule in the two channels.

Three-color 3D-STORM imaging was performed on targets labeled by Alexa Fluor 647, CF680, and CF568 via sequential imaging using 647-nm and 560-nm excitations. 647-nm excitation was first used to image Alexa Fluor 647 and CF680 as described above. Subsequently, 560-nm excitation was used to image CF568 through the reflected light path of the dichroic mirror.

---

**Figure S1.**
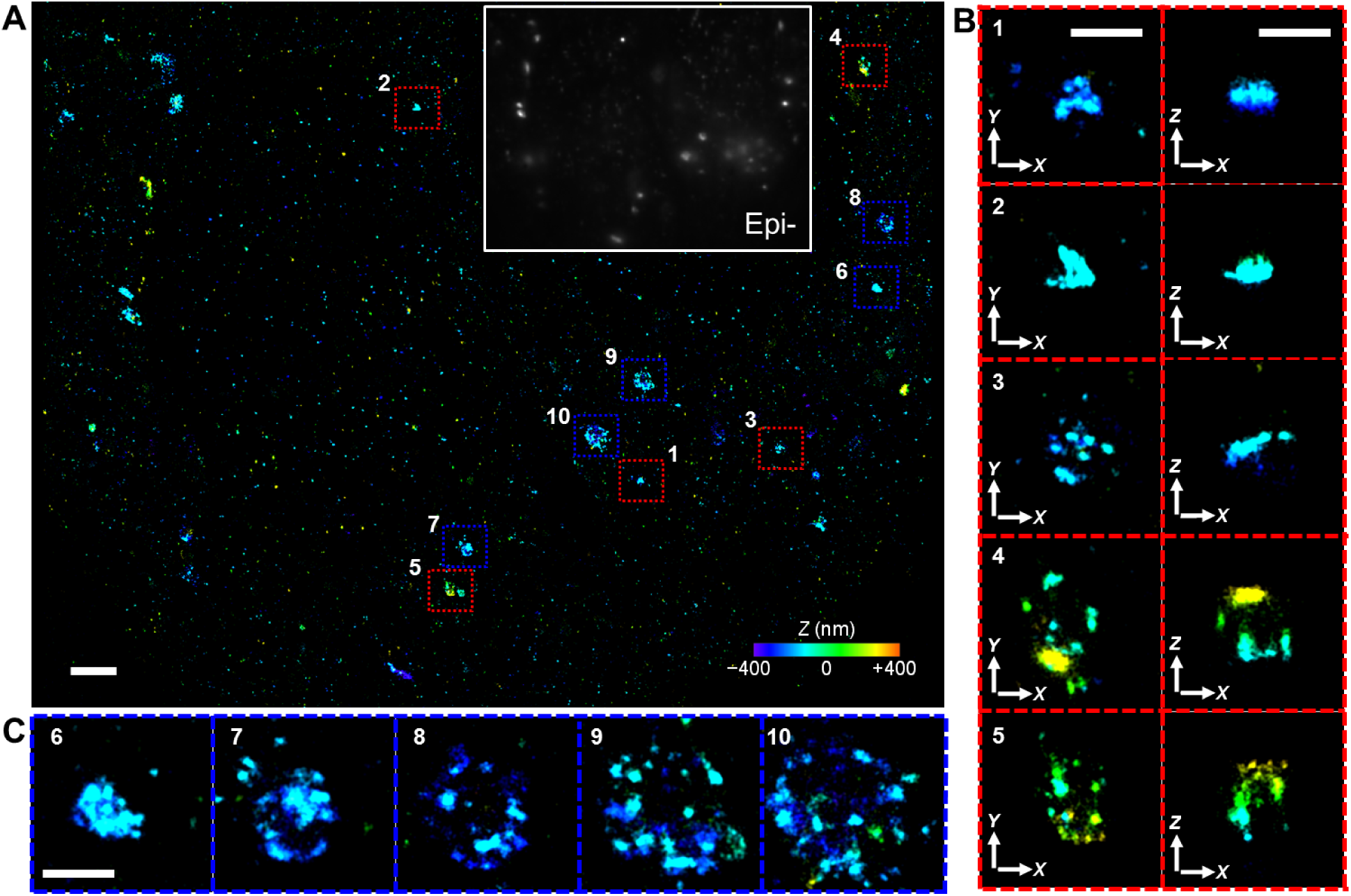
Additional 3D-STORM images of immunolabeled LC3 in starved U2OS cells. (A) A zoom-out view. Depth (*Z*) position is color-coded (color scale bar). Inset: diffraction-limited epi-fluorescence image of the same view. (B) Zoom-in of the structures enclosed by the red boxes 1-5 in (A). Left and right panels show projections in the X-Y and X-Z planes, respectively. (C) Zoom-in of the structures enclosed by the blue boxes 6-10 in (A). Scale bars: 2 μm (A); 500 nm (B,C).

**Figure S2.**
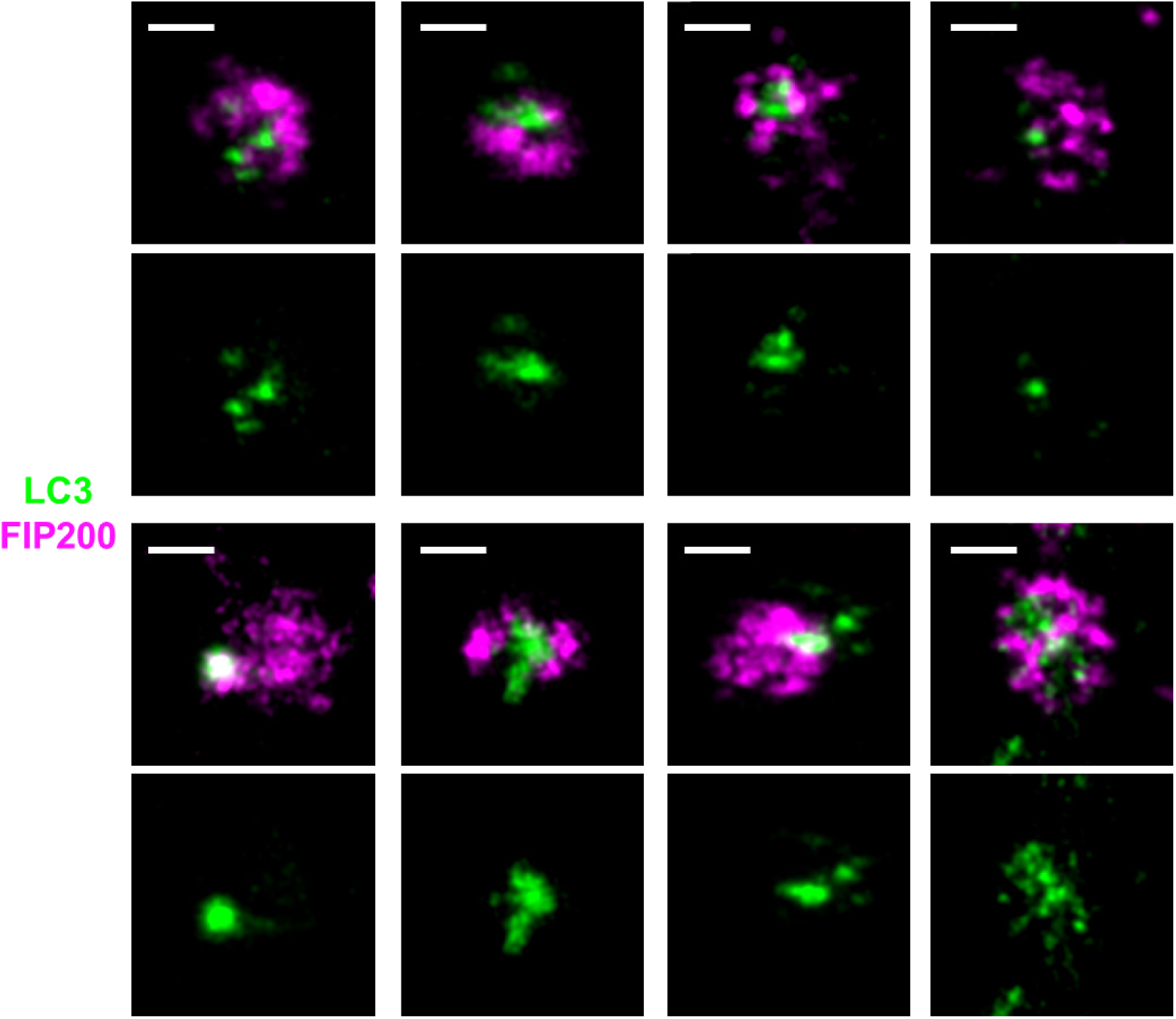
Additional representative early-stage cup structures, as identified by FIP200 (magenta) localization around a small (∼50 nm) LC3 (green) punctum. The average diameter of these cup structures provides an upper bound for their size prior to LC3 lipidation. Scale bars: 200 nm.

**Figure S3.**
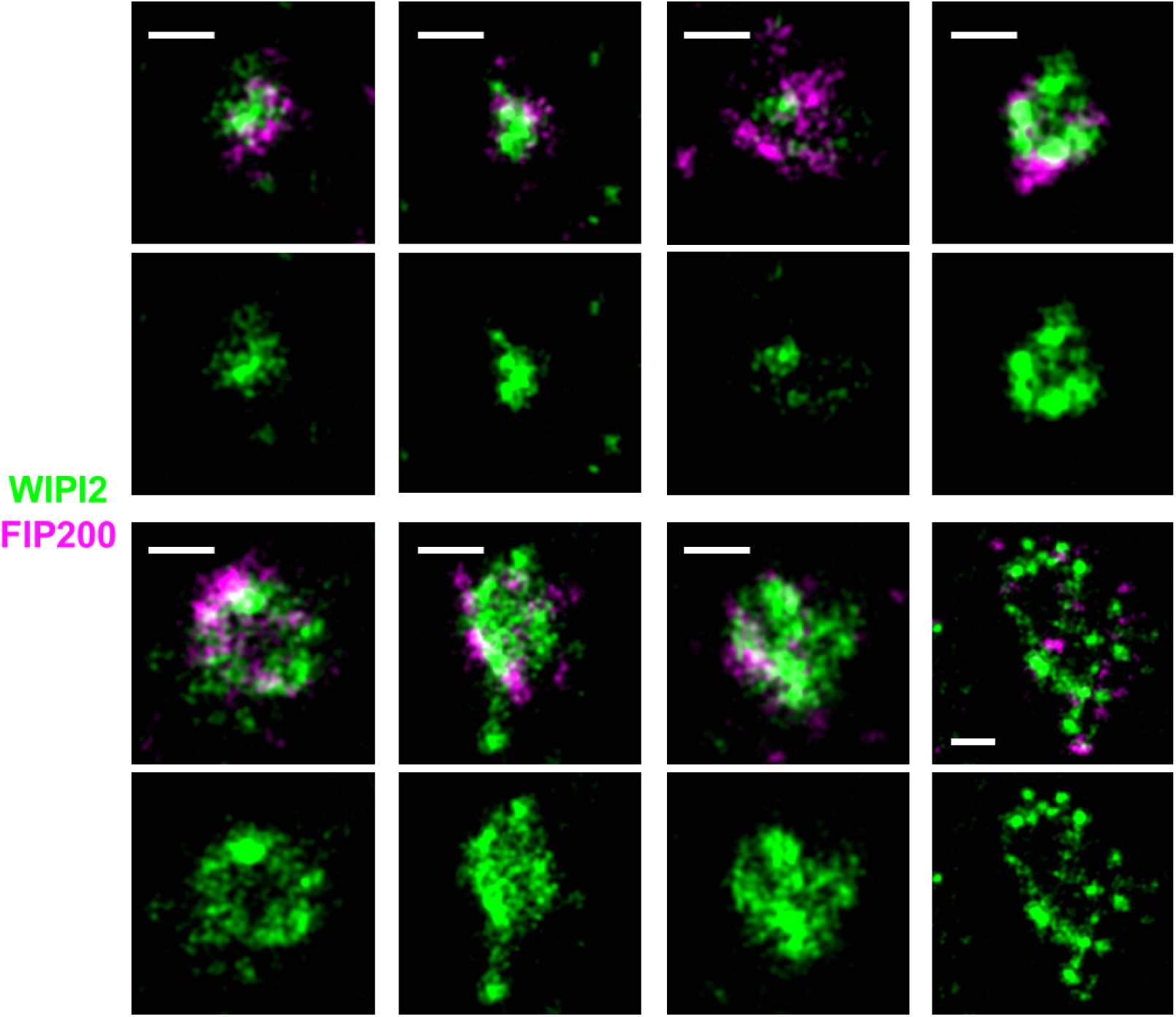
(A) Two-color 3D-STORM images of FIP200 (magenta) and WIPI2 (green). For each structure, the top and bottom panels show merged two-color images and the WIPI2 channel alone, respectively. Scale bars: 200 nm.

